# A Ca^2+^-ATPase Regulates E-cadherin Biogenesis and Epithelial-Mesenchymal Transition in Breast Cancer Cells

**DOI:** 10.1101/379586

**Authors:** Donna K. Dang, Monish Ram Makena, José P. Llongueras, Hari Prasad, Myungjun Ko, Manuj Bandral, Rajini Rao

**Affiliations:** Department of Physiology, The Johns Hopkins University School of Medicine, 725 N. Wolfe Street, Baltimore MD 21205

**Keywords:** *ATP2C2*, epithelial-mesenchymal transition, triple negative breast cancer, E-cadherin, Ca^2+^ signaling, store-independent Ca^2+^ entry (SICE), Hippo pathway

## Abstract

Progression of benign tumors to invasive, metastatic cancer is accompanied by the epithelial to mesenchymal transition (EMT), characterized by loss of the cell-adhesion protein E-cadherin. Although silencing mutations and transcriptional repression of the E-cadherin gene have been widely studied, not much is known about post-translational regulation of E-cadherin in tumors. We show that E-cadherin is tightly co-expressed with the secretory pathway Ca^2+^-ATPase isoform 2, SPCA2 (*ATP2C2*), in breast tumors. Loss of SPCA2 impairs surface expression of E-cadherin and elicits mesenchymal gene expression through disruption of cell adhesion in tumorspheres and downstream Hippo-YAP signaling. Conversely, ectopic expression of SPCA2 in triple negative breast cancer (TNBC) elevates baseline Ca^2+^ and YAP phosphorylation, enhances post-translational expression of E-cadherin, and suppresses mesenchymal gene expression. Thus, loss of SPCA2 phenocopies loss of E-cadherin in the Hippo signaling pathway and EMT-MET transitions, consistent with a functional role for SPCA2 in E-cadherin biogenesis. Furthermore, we show that SPCA2 suppresses invasive phenotypes, including cell migration *in vitro* and tumor metastasis *in vivo*. Based on these findings, we propose that SPCA2 functions as a key regulator of EMT and may be a potential therapeutic target for treatment of metastatic cancer.

**Implications:** Post-translational control of E-cadherin and the Hippo pathway by calcium signaling regulates epithelial mesenchymal transition in breast cancer cells.

## Introduction

Mortality from breast cancer is invariably due to tumor metastasis, which requires a key step known as epithelial to mesenchymal transition (EMT) during which cancer cells lose their polarity and cell-cell contacts, thereby acquiring the ability to migrate (1, 2). The process of EMT coincides with the loss of epithelial markers; the most critical of these is the Ca^2+^ binding cell adhesion protein E-cadherin (3, 4). As a structural component of the adherens junction in polarized epithelial cells, E-cadherin is a membrane-spanning protein that mediates homophilic cell-cell contacts through its extracellular domains and communicates these interactions via its intracellular domains to a network of signaling proteins to control cell polarity, organ size and epithelial integrity (5–7). Contact inhibition of growth is transmitted by E-cadherin to the Hippo signaling pathway, resulting in nuclear exclusion and inactivation of the transcriptional co-activator YAP (Yes-activated protein) and its paralog, TAZ (transcriptional activator with PDZ binding motif) (6, 8). Loss of Hippo signaling or overexpression of YAP/TAZ promotes metastatic phenotypes, including EMT (9). A better understanding of the molecular pathways that down-regulate E-cadherin and control the Hippo pathway may be pivotal to discovering new therapeutic targets for treatment of metastatic cancers.

E-cadherin expression is regulated at multiple levels, including genetic, epigenetic, transcriptional and post-translational. At the transcriptional level, *CDH1*, the gene encoding E-cadherin is repressed by a set of nuclear factors, including ZEB1 and SNAI2/SLUG that reprogram the cancer cell to acquire malignant invasive phenotypes (10). Expression of these factors in tumors inversely correlates with E-cadherin and prognosticates a more aggressive, invasive phenotype with poor clinical prognosis. In contrast to the wealth of information on *CDH1* regulation, much less is known of oncogenic control of E-cadherin biogenesis despite some evidence that post-translational down-regulation of E-cadherin precedes repression of transcription upon induction of EMT (11).

In this study, we uncover a novel functional link between E-cadherin and SPCA2, one of two secretory pathway Ca^2+^-ATPases that transport Ca^2+^ ions into the Golgi lumen for protein sorting, trafficking and quality control (12–14). Duplication of the SPCA gene (*ATP2C*) occurred during tetrapod evolution (15) and paved the way for acquisition of distinct functions. Although they have similar ion transport characteristics and appear redundant at first glance, we show that SPCA2, but not SPCA1, is an epithelial signature gene required for maximal cell surface expression of E-cadherin, and to maintain contact inhibition of EMT through the Hippo pathway. While both secretory pathway Ca^2+^ pumps supply the Golgi and downstream compartments with Ca^2+^, SPCA2 preferentially interacts with plasma membrane Ca^2+^ channels to elicit Ca^2+^ influx (16, 17). We show that ectopic expression of SPCA2 elevates baseline Ca^2+^ levels and YAP phosphorylation, resulting in suppression of mesenchymal phenotypes in triple receptor-negative breast cancer (TNBC) lines, including cell migration *in vitro* and tumor metastasis *in vivo*. Our findings reveal SPCA2 as a novel regulator of E-cadherin biogenesis and EMT in breast cancer cells, and a potential therapeutic target in the treatment of metastatic cancers.

## MATERIALS AND METHODS

### Hierarchical clustering

CCLE and TCGA invasive breast carcinoma project datasets were accessed through cBioPortal (http://www.cbioportal.org/) (18). Hierarchical clustering to analyze gene expression data was performed with statistiXL software and results were displayed as a dendrogram and correlation coefficient matrix heat map.

### Cell Culture

Cells used in this study were MCF7 (ATCC HTB-22), MDA-MB-231 (ATCC HTB-26), MCF10A (ATCC CRL-10317), SK-BR-3, BT-483, MDA-MB-415, AU-565 (gift from Dr. Saraswathi Sukumar, Johns Hopkins Medical School, MD). MDA-MB-231 tet-Ecad cells were a gift from the Barry Gumbiner, Seattle Children’s Hospital (6). SK-BR-3 and AU-565 were grown in RPMI 1640 medium containing antibiotic and 10% fetal bovine serum (FBS). MDA-MB-231 tet-Ecad and MCF7 cells were cultured in monolayer with DMEM containing antibiotic, antimycotic, and 10% serum. MCF10A monolayer cells were cultured in DMEM/F12 containing 5% horse serum, 20ng/ml EGF, 0.5mg/ml hydrocortisone, 100ng/ml cholera toxin, 10µg/ml insulin, and 1X penicillin/streptomycin. 50ng/ml of EGF (Sigma-Aldrich, cat#E9644) was added to the culture for 6 hours to induce EMT. Cells were cultured with 5% CO_2_ and at 37°C in a humidified incubator.

### Lentiviral Transfection

FUGW overexpression constructs and pLK0.1 shRNA and lentiviral construction of both SPCA isoforms was packaged and transfected according to previous methods using pCMV-Δ8.9 and PMDG at a ratio using 9:8:1 in HEK293T cells. A mixture of two shRNA constructs for SPCA2 gave the same results as individual constructs and have been previously validated, as reported (16, 19). Virus was collected after 48 hours and concentrated with Lenti-X Concentrator (Clontech, Cat#631231) (16). After 48 hours of transfection, cells were selected with puromycin at varying concentrations according to kill curves performed for each specific cell line (1 – 2.5 mg/mL). Experiments were performed within 5 passages to ensure maintained knockdown.

### cDNA synthesis & Quantitative PCR

1μg of RNA was collected and used for cDNA synthesis (Applied Biosystems, cat#4387406). The qPCR mastermix was made with EagleTaq Universal Mastermix (Roche, cat#07260296190), Taqman probe as specified, and 50ng of cDNA. Probes were as follows: Applied Biosystems; GAPDH (Hs02758991_g1), SPCA2 (Hs00939492_m1), SPCA1 (Hs00995930_m1), E-cadherin (Hs01023894_m1 or HS01023895_m1), N-cadherin (Hs00983056_m1), Zeb1 (Hs00232783_m1), Snail1 (Hs00195591_m1), and Vimentin (Hs00958111_m1).

### Western Blotting

35 – 50μg of lysates were utilized for protein analysis. GAPDH (Sigma-Aldrich, cat#G9295) was used as the loading control for Western blotting. E-cadherin (BD Transduction Laboratories, cat#610181), ZO-1 (H-300) (Santa Cruz Biotechnology, cat#sc-10804), pYAP (Ser127, Cell Signaling Technology #4911) and EGFR antibody (Abcam, cat# ab2430) was analyzed for total protein. Blots were analyzed using ImageJ.

### Immunofluorescence

Cells were cultured on glass coverslips and were rinsed with PBS and pre-extracted with 1X PHEM buffer, 8% sucrose and 0.025% saponin. Cells were fixed with 4% paraformaldehyde for 30 minutes and were rinsed and washed with PBS 3 times for 5 minutes each. After blocking in 1% BSA, cells were incubated with primary antibody overnight in 4°C. Cells were rinsed with 0.2% BSA 3 times for 5 minutes and were then incubated with a fluorescent secondary antibody in 1% BSA and 0.025% saponin buffer for 30 minutes at room temperature. Coverslips were washed and mounted onto slides with mounting media (Dako, cat#S3023). Immunofluorescent staining was analyzed using ImageJ.

### Flow Cytometry

Cells were rinsed with PBS and trypsinized and washed once with PBS. Resuspended cells were fixed in 4% paraformaldehyde at room temperature for 15 minutes and then washed 3 times in PBS. Fixed cells were permeabilized in a PBS buffer containing 0.1% saponin and 1% BSA for 15 minutes at room temperature. Cells were incubated with primary antibodies for 30 minutes at room temperature in the dark in the same saponin/BSA buffer and then washed 3 times. If needed, cells were incubated with secondary antibodies in the saponin/BSA buffer for 30 minutes in the dark and washed once. Cells were washed once and resuspended in a flow buffer (1mM EDTA, 0.1% Na^+^ azide, 1%BSA, PBS). Cells were run using BD FACSDiva on a BD LSR II flow cytometer and were analyzed with FlowJo. Antibodies used were 1:500 FITC Mouse Anti-E-cadherin (BD Biosciences, cat#612130), 1:500 DYKDDDDK Tag Polyclonal Antibody (ThermoFisher Scientific, cat#PA1-984B), 1:1000 Goat anti-Rabbit IgG (H+L) Cross-Adsorbed Secondary Antibody, Alexa Fluor 633 (ThermoFisher Scientific, cat#A-21070).

### Ca^2+^-dependent Fura2 Fluorescence

Live imaging of Ca^2+^ in MDA-MB-231 cells was performed as described (20). Fura2-AM (Invitrogen; 1μg/ml) was added to cells in imaging buffer (20 mM Hepes, 126 mM NaCl, 4.5 mM KCl, 2 mM MgCl_2_, 10 mM glucose at pH7.4). Cells were excited at 340 nm and 380 nm, and Fura emission was captured at 505 nm. To show store independent Ca^2+^ entry (SICE), cells were briefly washed in nominally Ca^2+^-free buffer followed by addition of Ca^2+^ (2mM) as described in the legend to Figure 5.

### Tumorsphere Suspension Assay

MCF7 cells (1000 cells) were seeded and cultured and in serum free suspension media (DMEM, 31% L-glutamine, 1% penicillin/streptomycin, 2% B27 Supplement, 20ng/ml EGF, and 20ng/ml FGFb) in 6-well culture plates according to the method from Iglesias et al. (21). Culture plates were treated with poly (2-hydroxethyl methacrylate) (Sigma-Aldrich, cat# P3932) prior to culturing to prevent adherence. Tumorspheres were counted and measured using ImageJ.

### YAP Localization

MCF-7 cells were infected with scramble “Control” or “shSPCA2” lentivirus for 48 hrs, and then selected with puromycin (2.5 mg/mL) for 48 hrs. Following, cells were seeded either sparse or confluent on coverslips, as previously described (22). Briefly, adherent cells were serum starved for 24 h before fixation and subsequent immunofluorescence. MCF7 cells, due to an aberrant mutation in PI3-K that alters YAP trafficking, were treated with wortmannin (10 mM) for 4 h to restore normal YAP trafficking prior to fixation (22). YAP intracellular localization was analyzed with anti-YAP antibody (63.7) (Santa Cruz Biotechnology, cat#SC-101199) using confocal microscopy. Where indicated, MDA-MB-231 cells expressing SPCA2R were treated with YAP inhibitor Verteporfin (10 μM; TOCRIS, Cat# 5305) for 24 hours, and investigated for changes in EMT markers.

### Cell Migration

#### Wound healing assay

MDA-MB-231 cells were plated in a 12-well culture plate. Upon reaching ~70% confluency, cells were maintained in 0.5% FBS containing DMEM media for 16 hours. Upon reaching confluency, using a pipette tip, horizontal scratches were made across the diameter of the well as described (23). The monolayer was washed couple of times with PBS to remove any floating cells generated by the scratching process, and fresh media (0.5% FBS containing DMEM) was added. Using an inverted microscope (Zeiss Axio observer), photos of each scratch were captured immediately at 2.5X magnification and used as a reference point for the 8 h, 16 h and 24 h time point to determine percentage of scratch coverage.

#### Boyden Chamber Assay

MDA-MB-231 cells were maintained in DMEM media without FBS for 4 hours. 3×10^4^ cells of each condition were plated in 100 μL DMEM media without FBS in 6.5 mm Transwell chambers with 8.0 μm Pore Polyester Membrane Insert (Corning, Cat# 3464), and transferred to a 24-well dish containing 500 μL fresh media (0.5% FBS containing DMEM). After 24 hours, cells were fixed with 4% paraformaldehyde for 30 min, 0.5% crystal violet (Sigma-Aldrich, Cat# V5265) was added for 1 hour, and washed with PBS. Using a microscope (Nikon, Eclipse TS100), photos of membrane insert were captured at 10X magnification. ImageJ software was used to quantify migration.

### Mouse Xenografts

Animal protocols described in this study were approved by the Institutional Care and Use Committee at the Johns Hopkins Medical Institutions. MDA-MB-231-luc-D3H2LN cell line expressing the luciferase gene was obtained from Cellosaurus. MDA-MB-231-luc-D3H2LN cells stably infected with SPCA2 overexpression or GFP containing virus. Cells were suspended in matrigel: DMEM media without FBS (1:1 volume). Six-to eight-week-old female NOD scid gamma (NSG) mice (The Jackson Laboratory) were anesthetized and approximately 0.7×10^6^ cells were injected into the mammary fat pad. Each group had 6 mice each. Animals were imaged every week to track tumor metastases using IVIS Lumina XR (Perkin-Elmer), after intraperitoneal injection of D-luciferin-K+ (Perkin-Elmer, 150 mg/kg). Metastasis was represented as mean total flux (p/sec) ± SEM, which was calculated using Living Image Software version 4.3.1 (Perkin-Elmer). Five weeks post injection, progression of metastasis between the GFP and SPCA2 overexpression groups was compared, and t-test was used to determine the significance. Mice weights were measured every week.

## Results

### SPCA2 is an epithelial marker tightly linked to E-cadherin expression

Gene expression correlations offer unique insight on disease pathophysiology, based on the premise that genes expressed together are likely to function together in a common pathway (24, 25). Both SPCA isoforms have been implicated in breast cancer oncogenesis and formation of breast microcalcifications linked to malignancy (16, 19, 26). Previously, we have evaluated subtype specific gene expression in breast tumors: whereas high SPCA1 is associated with basal subtypes, SPCA2 was found to be significantly elevated in luminal A/B and HER2+ subtypes (19). To further distinguish between their tumor expression pattern and gain insight into isoform-specific functions, we compared the expression of SPCA1 (*ATP2C1*) and SPCA2 (*ATP2C2*) genes in breast cancer patient specimens from TCGA (n=526) with a “benchmark” gene set, consisting of 18 epithelial signature genes (ESG) and 13 mesenchymal signature genes (MSG), described previously (24). As expected, we observed a sharp dichotomy between ESG and MSG, with the epithelial and mesenchymal genes separating as two tightly distinct and mutually correlated clusters (Fig. 1A). SPCA1 and SPCA2 showed differential clustering with MSG and ESG, respectively, despite their high identity (64%) in amino acid sequence and similar Ca^2+^ transport characteristics (27, 28). Unexpectedly, we observed a strong co-expression correlation between SPCA2 and E-cadherin (Fig. 1A; Fig. S1A, Pearson coefficient 0.44; *p* = 2.52 × 10^−26^). In contrast, there was no significant correlation between SPCA1 and E-cadherin expression (Fig. S1B).

**Figure 1.**
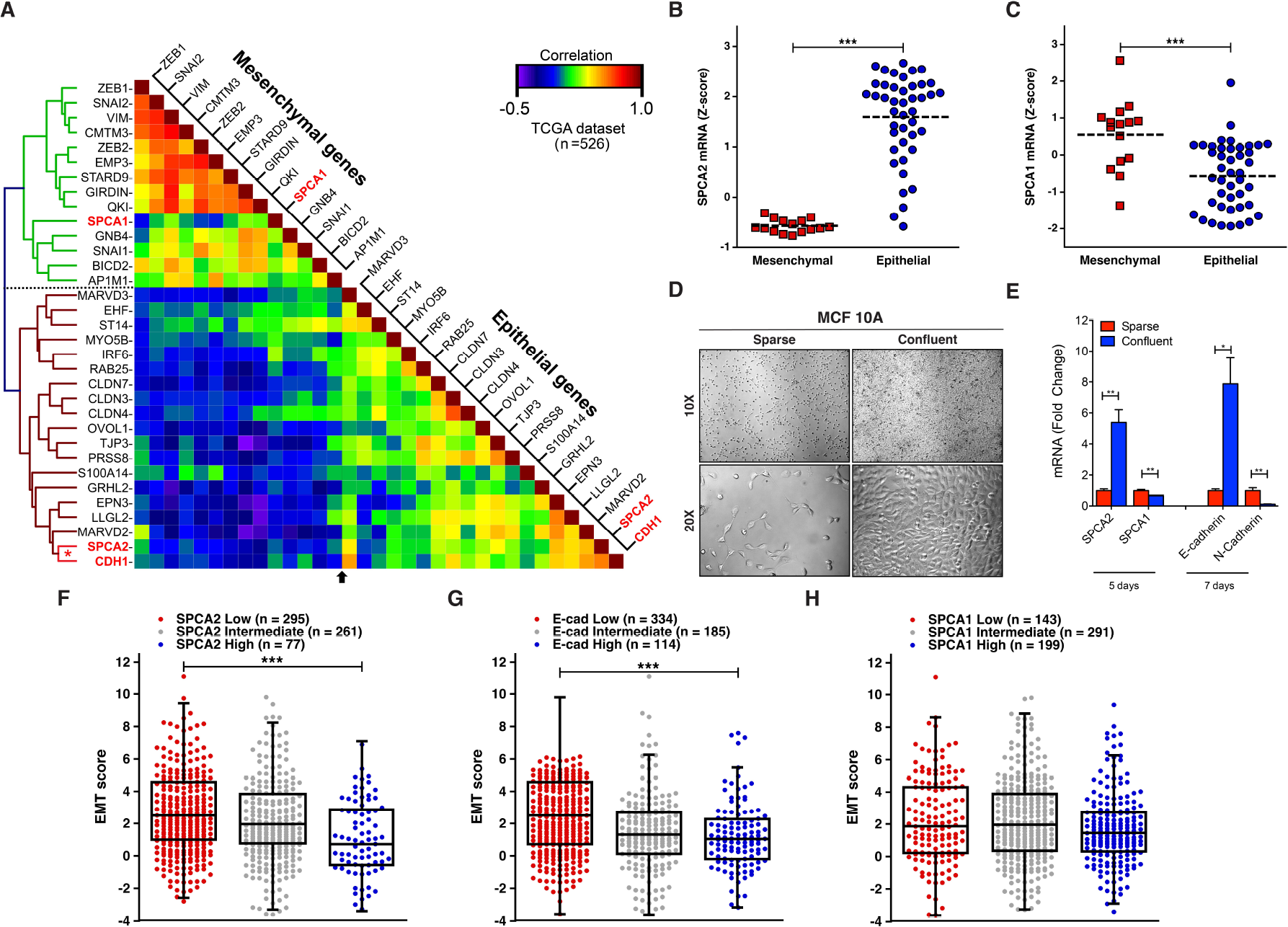
SPCA2 is an epithelial marker co-expressed with E-cadherin. (A) Hierarchical clustering and heat map of the correlation coefficient matrix of SPCA isoforms with ESG and MSG in breast cancer samples from TCGA (low correlation, blue; high correlation, red; n=526) (49). SPCA1 and SPCA2 cluster with MSG and ESG, respectively. Note the strong correlation of SPCA2 with *CDH1*(red asterisk). (B-C) Breast cancer cell lines from CCLE grouped as mesenchymal or epithelial to show Z-scores for SPCA2 (B) and SPCA1 (C) mRNA (24, 50). Note, significant up-regulation of SPCA2 and conversely, significant down-regulation of SPCA1 in epithelial-like breast cancer cell lines. ***p<0.001, Student’s t-test. (D) MCF10A cells were cultured in both sparse and confluent conditions and DIC images were taken with 10X and 20X objectives on a Zeiss Axio Observer A.1 microscope. (E) qPCR analysis of SPCA2, SPCA1, E-cadherin and N-cadherin expression in cDNA extracted from sparse cells or after 5 or 7 days post-confluence. ***p<0.001, **p<0.01, two-tailed Welch’s t-test, n=3. (F-H) Box and whisker with scatter plots of the EMT score across 633 breast cancer samples from the TCGA dataset grouped into low, intermediate and high expression categories for SPCA2 (F), E-cadherin (G) and SPCA1 (H) (***p<0.001) (30). Notably, tumors with low SPCA2 or E-cadherin expression had significantly higher EMT score than those with high SPCA2 expression. See Fig. S1.

To corroborate these findings, we evaluated SPCA gene expression in breast cancer cell lines of the Broad Institute’s Cancer Cell Line Encyclopedia (CCLE), previously classified into epithelial and mesenchymal subtypes based on their consensus signatures (24). In agreement with Fig. 1A, there was up-regulation (1.89-fold change) of SPCA2 (Fig. 1B) and conversely, down-regulation (0.61-fold change) of SPCA1 (Fig. 1C) in epithelial-like breast cancer cell lines based on Z-score which shows the number of standard deviations above or below the mean (Z-score=0). To determine if these expression patterns extended to non-malignant mammary epithelial cells we examined cell density related changes in gene expression in sparse and confluent cultures of MCF10A. Unlike cancer cells, sparsely seeded MCF10A cells ceased to proliferate upon reaching confluence (Fig. 1D), due to contact inhibition and mesenchymal to epithelial transition (MET). Within 5 days post-confluence, we observed up-regulation of SPCA2 by 5-fold in MCF10A cultures, whereas SPCA1 levels decreased by 2-fold (Fig. 1E). These changes preceded the so-called “cadherin switch” that accompanies MET, characterized by increase in E-cadherin expression and reciprocal decrease of N-cadherin which we observed at 7-days post-confluence (Fig. 1E). These findings were replicated in the breast cancer cell line MDA-MB-231 (Fig. S1C-F). Similarly, cell-density dependent up regulation of SPCA2, but not SPCA1, has been observed in the human colon cancer line HCT116 (29). This distinctive transcriptional signature of Golgi calcium pumps suggests that the two isoforms play different cellular roles related to EMT-MET transitions.

Tumor EMT scores quantitatively capture the molecular features of cancer cells exhibited during EMT (25). Tumors with high EMT score have high potential for invasion and metastasis. We analyzed 633 breast cancer samples of all subtypes from TCGA to probe the link between EMT score and SPCA2 gene expression (30). Using Z-score of ±0.5 as cut-off for SPCA2 expression, breast cancer samples were stratified into low, intermediate and high expression categories. Tumors with low SPCA2 expression had significantly higher EMT score than those with high SPCA2 expression (Fig. 1F, p<0.001; Fig. S1H, corr. −0.18, p=0.0009). This correlation was similar in tumors sorted for E-cadherin expression (Fig. 1G, p<0.001; Fig. S1G corr. −0.25, p<0.0001). No such correlation was observed for SPCA1 (Fig. 1H, Fig. S1I). Taken together, the robust correlation in gene expression between SPCA2 and E-cadherin point to a potential EMT-related functional link in breast cancer cells.

### SPCA2 regulates E-cadherin biogenesis

Within the secretory pathway, Ca^2+^ is important for sorting and trafficking of cell surface proteins (31, 32). Therefore, we asked if SPCA2 was required for E-cadherin biogenesis using two complementary approaches: gene knockdown in the high-SPCA2 expressing breast cancer cell line MCF7 and ectopic gene expression in the low-SPCA2 expressing line MDA-MB-231. Previously, we showed high levels of SPCA2 in ER+, luminal subtype MCF7 cell line, which we confirmed in this study (see ahead, Fig 3A). Targeted knockdown of SPCA2 in MCF7 cells did not alter E-cadherin mRNA (Fig. 2A). However, total E-cadherin protein was significantly reduced (53 ± 7.6% of control; *p* < 0.05; Fig. 2B-C), suggesting an upstream requirement for SPCA2 in E-cadherin expression. These observations were replicated in MCF10A cells (Fig. S2A-D). In contrast, there was no significant change in E-cadherin protein expression upon SPCA1 knockdown (Fig. S2E-F) as previously reported in keratinocytes (30). Furthermore, there was no reduction of epidermal growth factor receptor (EGFR), a surface protein not found at the adherens junction (Fig. S2G-H), or of zona occludens-1 (ZO-1), a tight junction protein, in shSPCA2-treated cells (Fig. S2I-J). Thus, the selective defect in E-cadherin biogenesis points to a functional link with SPCA2 and is consistent with the results of the co-expression analysis (Fig. 1).

**Figure 2.**
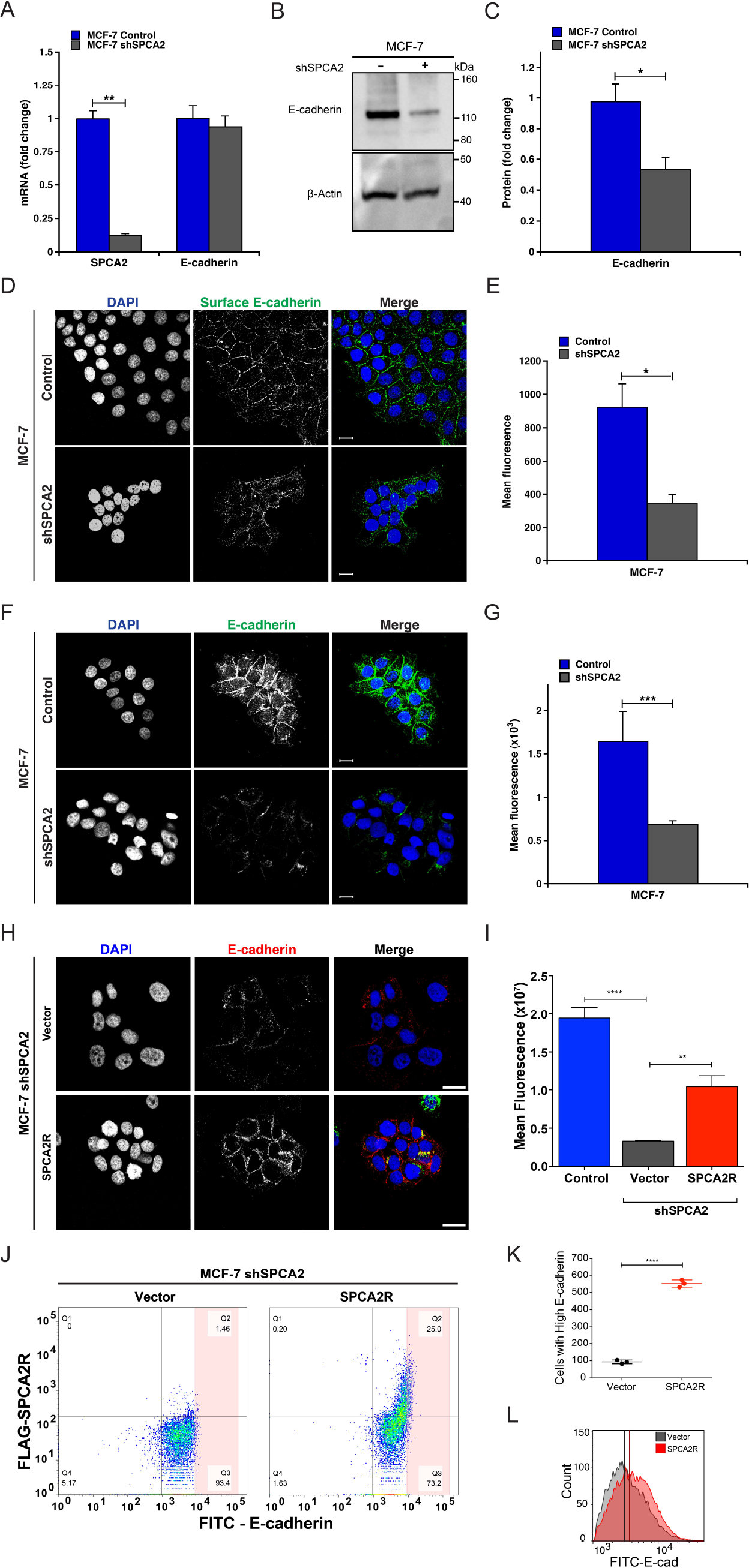
Loss of SPCA2 impairs E-cadherin biogenesis. (A) Knockdown of SPCA2 in MCF7 cells was confirmed by qPCR. There was no change in E-cadherin transcript. **p<0.01, Student’s t-test. (B) Western blot of cell lysates probed with antibody against E-cadherin and β-Actin. (C) E-cadherin protein expression from densitometric analysis of Western blots, normalized against β-Actin. *p<0.05, Student’s t-test, n=3. (D) Immunofluorescence of surface E-cadherin in non-permeabilized MCF7 cells treated with both control and shSPCA2. Note that the antibody recognizes an extracellular epitope. (E) Total mean fluorescence was quantified. *p<0.05, Student’s t-test, control n=276, shSPCA2 n=47. (F) Immunofluorescence of E-cadherin in permeabilized MCF7 cells treated with both control and shSPCA2. Note that the antibody recognizes a cytoplasmic epitope (G) Total mean fluorescence from (F) was quantified. ***p<0.001, Student’s t-test, control n=91, shSPCA2 n=44. (H) Immunofluorescence of total E-cadherin in permeabilized MCF7 shSPCA2 cells expressing vector control, or FLAG-tagged SPCA2R (shRNA-resistant SPCA2), with quantification (I). *p<0.05, **p<0.01, Student’s t-test, vector n=34, SPCA2R n=44. Scale bars are 20 μm. (J) Flow cytometry analysis of MCF7 shSPCA2 cells transfected with empty vector or SPCA2R showing percentages of cells in each quadrant (Q1-Q4). (K) Number of cells with high E-cadherin fluorescence found in the pink shaded area, representing 0.78% and 5.07% of total cells in Vector or SPCA2R transfected cells, respectively. (L) Histogram showing shift in total E-cadherin fluorescence in shSPCA2 cells expressing SPCA2R. *p<0.05, ***p<0.001, Student’s t-test, n=3 for each condition. See SI Appendix, Fig. S2.

**Figure 3.**
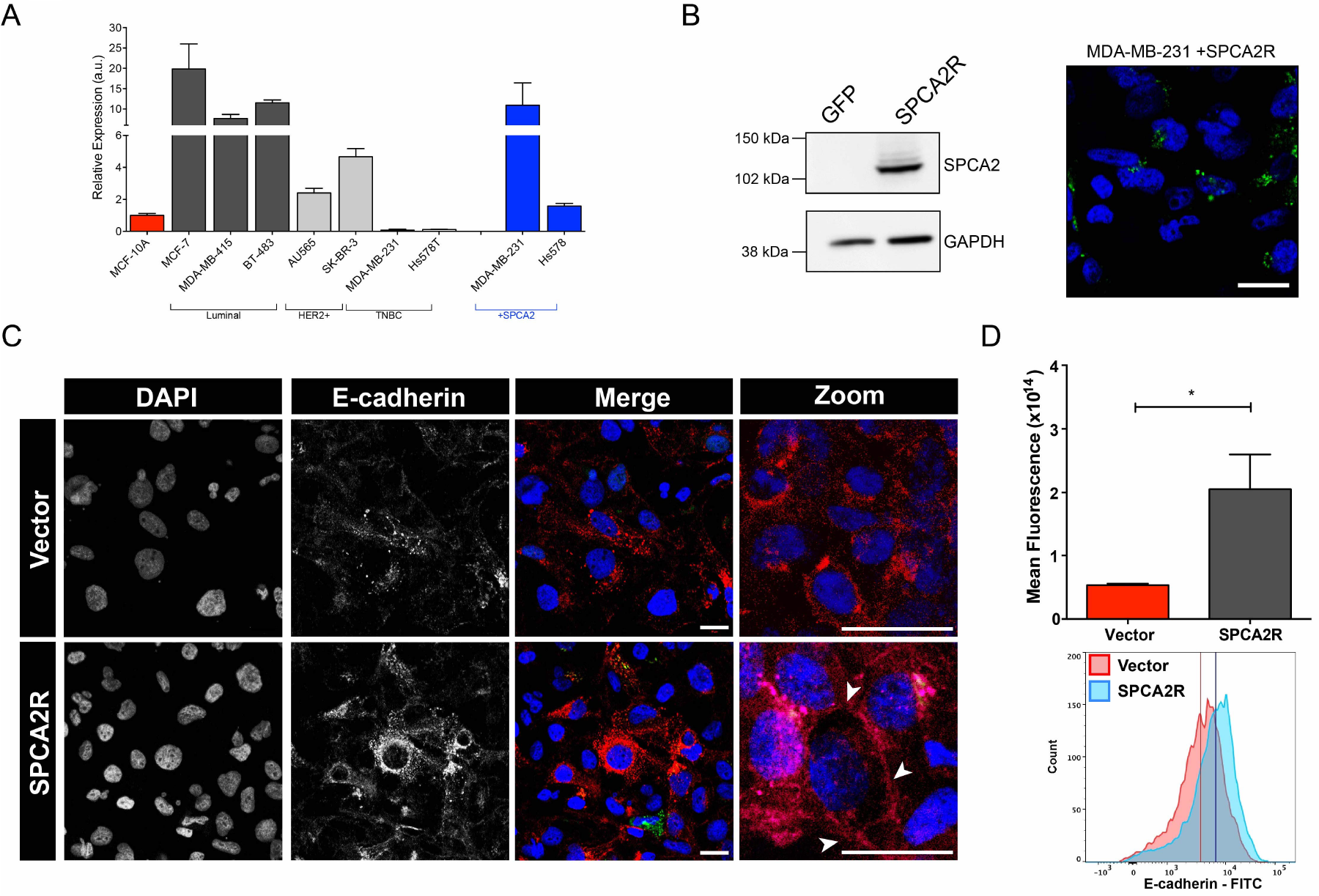
SPCA2 increases E-cadherin expression in TNBC cells. (A) A panel of breast cancer cell lines classified as luminal, HER2+ or TNBC subtypes as indicated, was evaluated for SPCA2 expression by qPCR. Results are normalized to MCF-10A. SPCA2 transcripts were lowest in two TNBC lines, MDA-MB-231 and Hs578T, and could be significantly increased by lentiviral-mediated transfection of SPCA2R. (B) SPCA2R expression in transfected MDA-MB-231 cells was confirmed by Western blotting (left) and confocal immunofluorescence microscopy (right) using anti-FLAG antibody. (C) MDA-MB-231 Tet-Ecadherin cells expressing either vector or SPCA2R were treated with 2μg/ml of doxycycline, stained for E-cadherin and imaged using confocal microscopy. (D) Normalized fluorescence values for each condition (top) and histograms of cells separated by flow cytometry (bottom). *p<0.05, Student’s t-test, vector n=3, SPCA2R n=3. Flow cytometry histogram analysis of 3 biological replicates of each condition. Vector n=9468 cells, SPCA2R n=9258 cells. See Fig. S3.

Treatment of non-permeabilized MCF7 cells with antibody directed against an external epitope of E-cadherin, revealed a significant reduction in surface expression of E-cadherin in SPCA2-depleted cells (Fig. 2D-E). Total E-cadherin level in SPCA2-depleted MCF7 cells, as determined by quantitative immunofluorescence microscopy (Fig. 2F-G), was also diminished but could be largely restored by transfection of a FLAG-tagged and shRNA-resistant construct SPCA2R, relative to vector-transfected or untreated controls (Figure 2H-I). This was confirmed by flow cytometry analysis: SPCA2R increased E-cadherin expression in knockdown cells relative to the empty vector control, as seen by forward scatter in the upper right quadrant (Figure 2J), increased number of cells with high E-cadherin (Figure 2K), and shift in the histogram of cell population with E-cadherin fluorescence (Figure 2L).

Since loss of SPCA2 was sufficient to down-regulate expression of E-cadherin in MCF7, an epithelial-subtype tumor cell line, we asked if ectopic expression of SPCA2 in mesenchymal-subtype tumor cells conferred a reciprocal increase in E-cadherin expression. Evaluation of a panel of established breast cancer cell lines revealed very low SPCA2 transcript levels in the TNBC cell lines MDA-MB-231 and Hs578T relative to MCF10A (Fig. 3A); in contrast, SPCA1 was expressed at levels similar to MCF10A or significantly higher in these cell lines (Fig. S3). Lentiviral-mediated transfection with SPCA2R in the TNBC lines increased transcripts to levels approaching endogenous SPCA2 expression in MCF7 cells, although this was cell-type specific (Fig. 3A, see also Fig. S6A,G). We confirmed ectopic expression of FLAG-tagged SPCA2R in MDA-MB-231 cells by Western blotting, and localization to intracellular vesicular compartments by confocal microscopy (Fig. 3B). By following doxycycline-induced E-cadherin protein in Tet-Ecad-MDA-MB-231 cells, we focused only on post-translational effects of SPCA2. E-cadherin levels were significantly increased by SPCA2R relative to vector control, as shown by both confocal microscopy and FACS analysis (Figure 3C-D) of immunofluorescent cells. Although the bulk of E-cadherin remained within the cell, suggesting that additional factors may be necessary to restore efficient surface trafficking of E-cadherin in TNBC cells, some E-cadherin reached the cell boundary as seen in the enlarged images (arrow heads, Fig 3C). Together, these data point to a hitherto unrecognized, post-translational role for SPCA2 in E-cadherin expression.

### SPCA2 phenocopies E-cadherin in tumorsphere formation

Next, we asked if SPCA2 regulated cellular phenotypes downstream from E-cadherin. The ability of breast cancer cells to form free-floating spheroids known as tumorspheres, in suspension culture has been shown to depend on surface expression of E-cadherin (21, 33). Thus, knockdown of E-cadherin in MCF7 abolished tumorsphere formation (21). Therefore, we asked if SPCA2-depletion phenocopied loss of E-cadherin in tumorsphere formation.

Depletion of SPCA2 in MCF7 cells (confirmed in Fig. 2A) resulted in nearly complete loss of tumorspheres, strongly attenuating both number and size relative to scrambled shRNA control (Fig. 4A). These results confirm and extend our previous finding that SPCA2 knockdown significantly inhibits formation of mammospheres derived from normal mammary epithelial cells embedded in extracellular matrix (20). In contrast, SPCA1 knockdown had no significant effect on tumorsphere formation (Fig. 4A). Surface expression of E-cadherin and tumorsphere formation could be rescued by re-introduction of SPCA2R in shSPCA2 treated MCF7 cells (Fig. 4B-D). Thus, tumorsphere formation closely parallels expression of both SPCA2 and E-cadherin (21).

**Figure 4.**
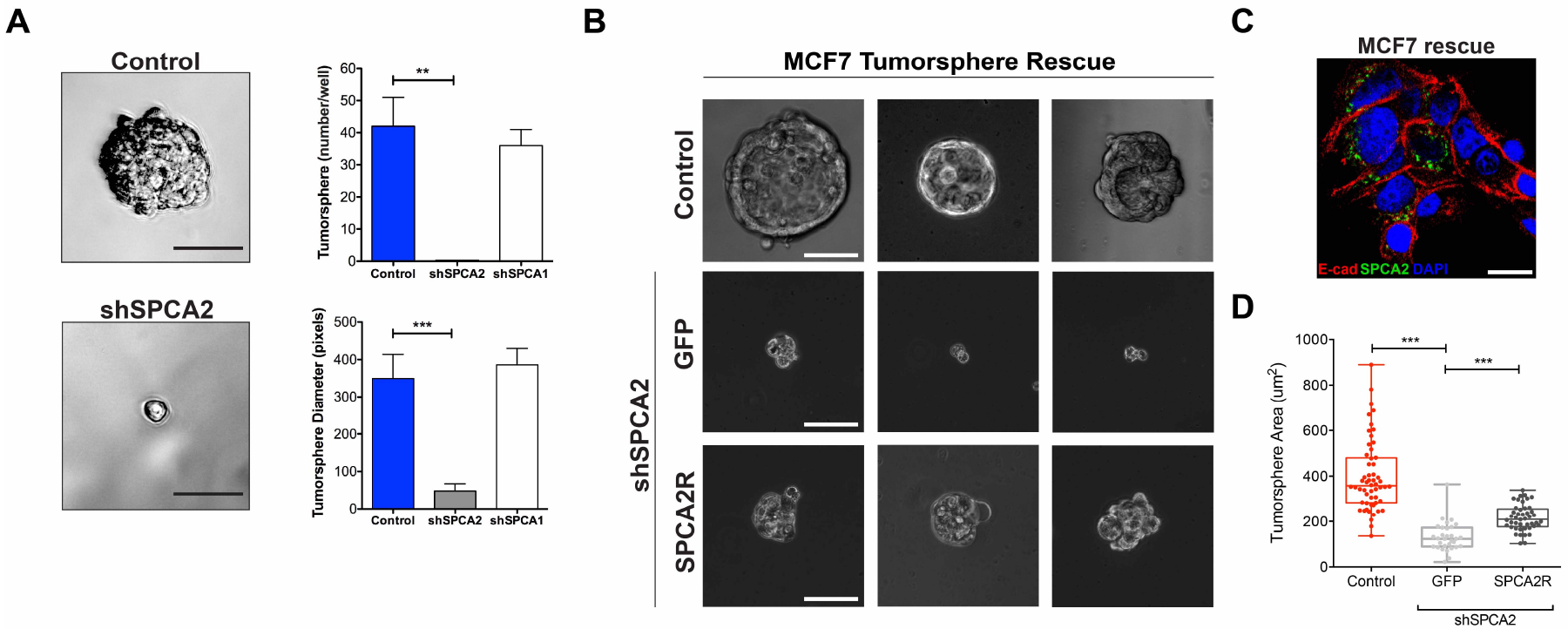
SPCA2 is required for tumorsphere formation. MCF-7 cells treated with scramble shRNA (control), shSPCA1 or shSPCA2 constructs were cultured to form tumorspheres. (A) Representative images of control and shSPCA2 cells are shown respectively in the upper and lower panels. Scale bars are 50μm. The numbers of tumorspheres (upper panel) and their diameters (lower panel) were quantified. **p<0.01, ***p<0.001, Student’s t-test, n=3. (B) Three representative images of MCF7 cells (untreated control), or shSPCA2-transfected cells rescued with empty vector (GFP) or SPCA2R, and cultured as tumorspheres. Scale bars are 50μm. (C) Immunofluorescence image showing that FLAG-tagged SPCA2R (green) effectively rescues surface E-cadherin expression (red) in permeabilized MCF7 shSPCA2 cells grown in 2-D culture. (D) Tumorsphere area from the experiment in (B) was quantified. ***p<0.001, Student’s t-test; for tumorsphere area, n=31 for vector, n=45 for SPCA2R. See Fig. S4.

**Figure 5.**
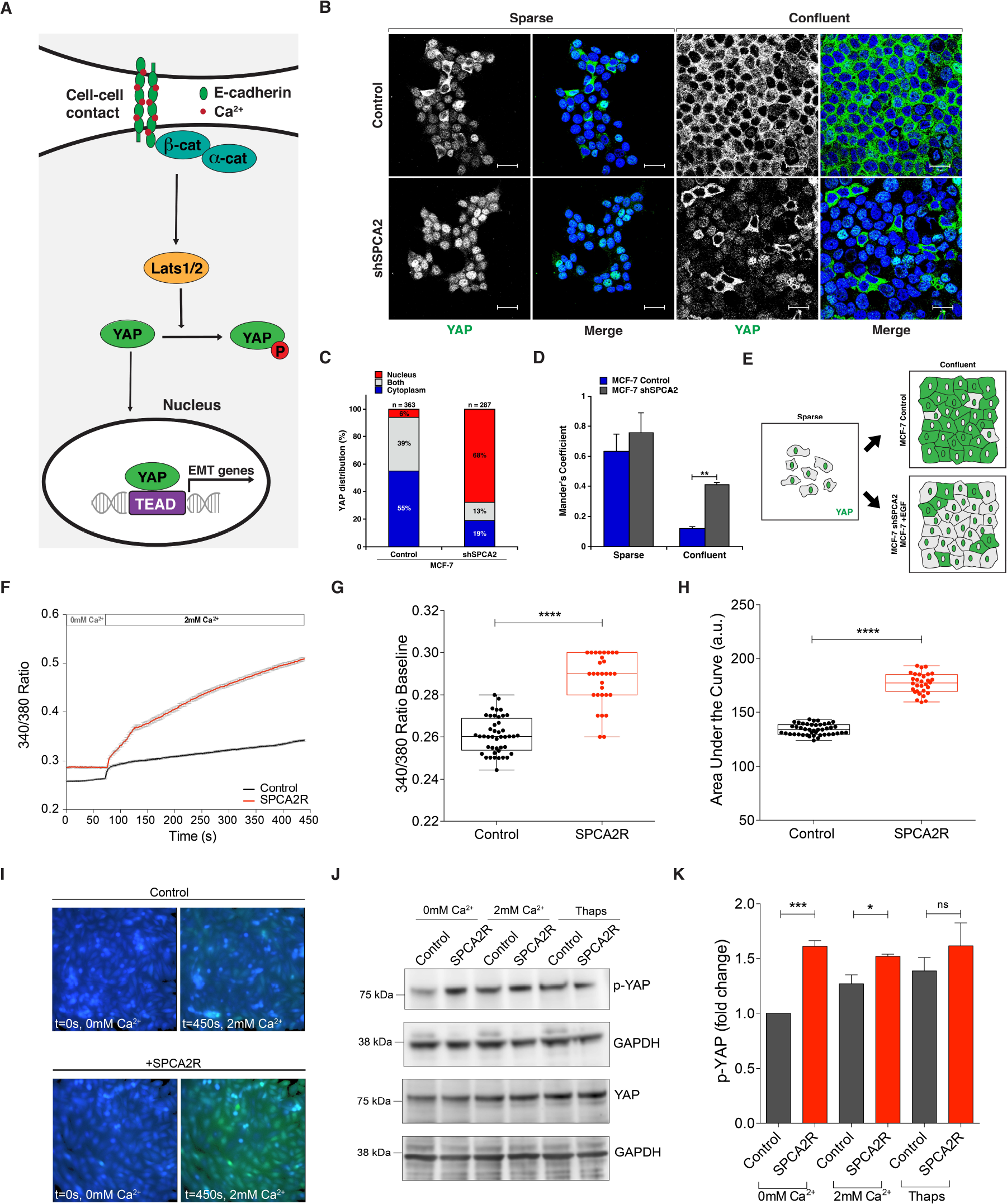
SPCA2 functions upstream of Hippo-YAP signaling. (A) Schematic showing the role of Hippo-YAP signaling in control of EMT. Cell-cell contacts mediated by E-cadherin maintain nuclear exclusion of YAP, suppressing expression of EMT genes. (B) Immunofluorescence staining of YAP in MCF7 cells treated with scramble (control) or shSPCA2 and cultured under sparse or confluent conditions. Note the distribution of YAP localization between nuclei (stained with DAPI, blue) and cytoplasm. (C) Subcellular YAP distribution in MCF7 cells grown to confluence and stained as in (B). (D) Colocalization of YAP with DAPI staining in confluent MCF7 cells using Manders’ coefficient. **p<0.01, Student’s t-test. (E) Schematic of YAP localization in response to cell density, EGF treatment and SPCA2 knockdown. (F) Live cell Ca^2+^ imaging traces using Fura2-AM treated MDA-MB-231 cells with or without SPCA2R (Control n=44 cells, SPCA2R n=30 cells). Baseline readings in calcium-free conditions were established for 50 cycles (72 s) and then cells were recorded under 2mM Ca^2+^ conditions for 250 cycles (6 minutes and 20 s). Baseline ratios (G) and the area under the curve (H) were plotted between both Control (n=44) and SPCA2R (n=30) samples. ****p < 0.0001, Student’s t-test. Images representing time points at 0 and 450 s of Control cells and cells expressing SPCA2R (I) showing calcium influx. (J) Western blot of cell lysates showing phospho-YAP and total YAP protein in Ca^2+^ free media and following addition of Ca^2+^ (2mM) or thapsigargin (1 μM). (L) Quantification of pYAP after normalization to GAPDH levels. Fold change (n=3 independent replicates; Student’s t-test) is shown. See Supplemental Movies 1 and 2.

Tumorsphere formation is commonly used to indicate the presence of cancer stem cells. Interestingly, despite the inability to form tumorspheres, depletion or overexpression of SPCA2 in MCF7 and MDA-MB-231 respectively, resulted in corresponding increase and decrease in the expression of genes associated with cancer stem cells, including *OCT4*, *NANOG* and *SOX2* (Fig. S4). These findings suggested that SPCA2 is an upstream regulator of stemness and post EMT-phenotypes.

### SPCA2 is an upstream regulator of YAP and Hippo pathway signaling

E-cadherin is required for cell-adhesion and contact inhibition of proliferation (34). By activating the Hippo tumor suppressor signaling pathway, E-cadherin controls phosphorylation, inactivation and nuclear exclusion of YAP, resulting in inhibition of cell proliferation in normal human mammary epithelial and breast cancer cells (Fig. 5A) (6, 8). Given the early post-confluency induction of SPCA2 gene expression ahead of the cadherin switch (Fig. 1E), together with a role for SPCA2 in efficient surface expression of E-cadherin (Fig. 2), we postulated that SPCA2 is an upstream regulator of the Hippo pathway. We tested this hypothesis both by knockdown of SPCA2 in MCF7 cells and by introduction of SPCA2 in MDA-MB-231 cells. First, MCF7 cells were cultured under sparse and confluent conditions and immunostained for YAP localization. As previously described for control cells, nuclear occupancy of YAP in sparsely growing cells largely reverted to the cytoplasm upon reaching confluency (Fig. 5B, top panel) (6). In contrast, in cells depleted of SPCA2, YAP remained predominantly in the nucleus under both sparse and confluent culture conditions (Fig. 5B, bottom panel; Fig. 5C-E). Again, loss of SPCA2 phenocopies loss of E-cadherin and downstream components of the Hippo signaling pathway (6), consistent with a functional role for SPCA2 in E-cadherin biogenesis.

One surprising finding was the selective requirement for SPCA2, but not SPCA1, for E-cadherin expression. Previously, we showed an isoform-selective role for SPCA2 in activating the Ca^2+^ channel Orai1 to elicit robust Ca^2+^ influx in MCF7 cells (16). Recent findings in a glioblastoma model have revealed that Ca^2+^ is a crucial intracellular cue that regulates the Hippo pathway by activation of a protein kinase C (PKC) beta II that leads to YAP phosphorylation and nuclear exclusion via upstream kinases Lats1/2 (35). Here, we used ectopic SPCA2 expression in MDA-MB-231 cells to elevate resting levels of Ca^2+^ (Fig. 5F-G). Addition of Ca^2+^ (2mM) to cells briefly incubated in Ca^2+^-free medium elicited immediate and sustained store-independent Ca^2+^ entry (SICE) that was significantly larger in SPCA2-transfected cells (Fig. 5F, H, I and Supplemental Movies 1 and 2). In parallel, we observed increased YAP phosphorylation under baseline conditions in SPCA2-transfected MDA-MB-231 cells, relative to the vector control (Fig. 5J-K). YAP phosphorylation further increased following Ca^2+^ influx from the media (addition of 2 mM Ca^2+^) or internal stores (1 μM thapsigargin treatment; Fig 5J-K), as expected. Thus, store-independent Ca^2+^ influx elicited by SPCA2 expression increases YAP phosphorylation associated with nuclear exclusion, extending the emerging observations from glioblastoma cells (35), and maintains Hippo pathway signaling downstream of E-cadherin in breast cancer cells.

### SPCA2 antagonizes epithelial mesenchymal transition

Nuclear YAP, and the related co-activator TAZ, interact with the TEAD family of transcriptional activators to elicit EMT by turning on mesenchymal markers such as N-cadherin and vimentin and promote malignant transformation (36). Furthermore, YAP/TAZ interact with multiple members of the EMT-associated family of transcription factors, including SNAI1, SLUG/SNAI2 and ZEB1 (37). A proteomic analysis of MCF7 cells showed that E-cadherin knockdown resulted in elevated expression of the EMT-associated transcription factors SLUG/SNAI2 and ZEB1, downstream of YAP (38). Interestingly, expression of YAP/TAZ was negatively correlated with SPCA2 expression in patient breast cancer tissue from all cancer subtypes (Fig. S5 A-B; *p* = 3.23 × 10^−7^, n = 295; data from GSE31448 microarray dataset (39)). Specifically, high YAP/TAZ expression is correlated with SPCA2 down-regulation in basal-like breast cancer (Fig. S5A). No correlation was observed between YAP/TAZ expression and that of SPCA1 (Fig. S5C; *p* = 0.178, n = 295). To test the hypothesis that SPCA2 regulated EMT-MET transitions (Fig. 6A), we elicited EMT in MCF7 cells by treatment with EGF. Although SPCA1 transcript levels remained unchanged, SPCA2 levels decreased substantially upon EGF treatment (Fig. 6B). Confirming the induction of EMT, cells treated with EGF showed enhanced expression of most EMT-associated genes including ZEB1, N-cadherin, SLUG/SNAI2 and Fibronectin, as expected (Fig. 6C-G, blue bars). Thus, SPCA2 gene expression correlates with EMT.

**Figure 6.**
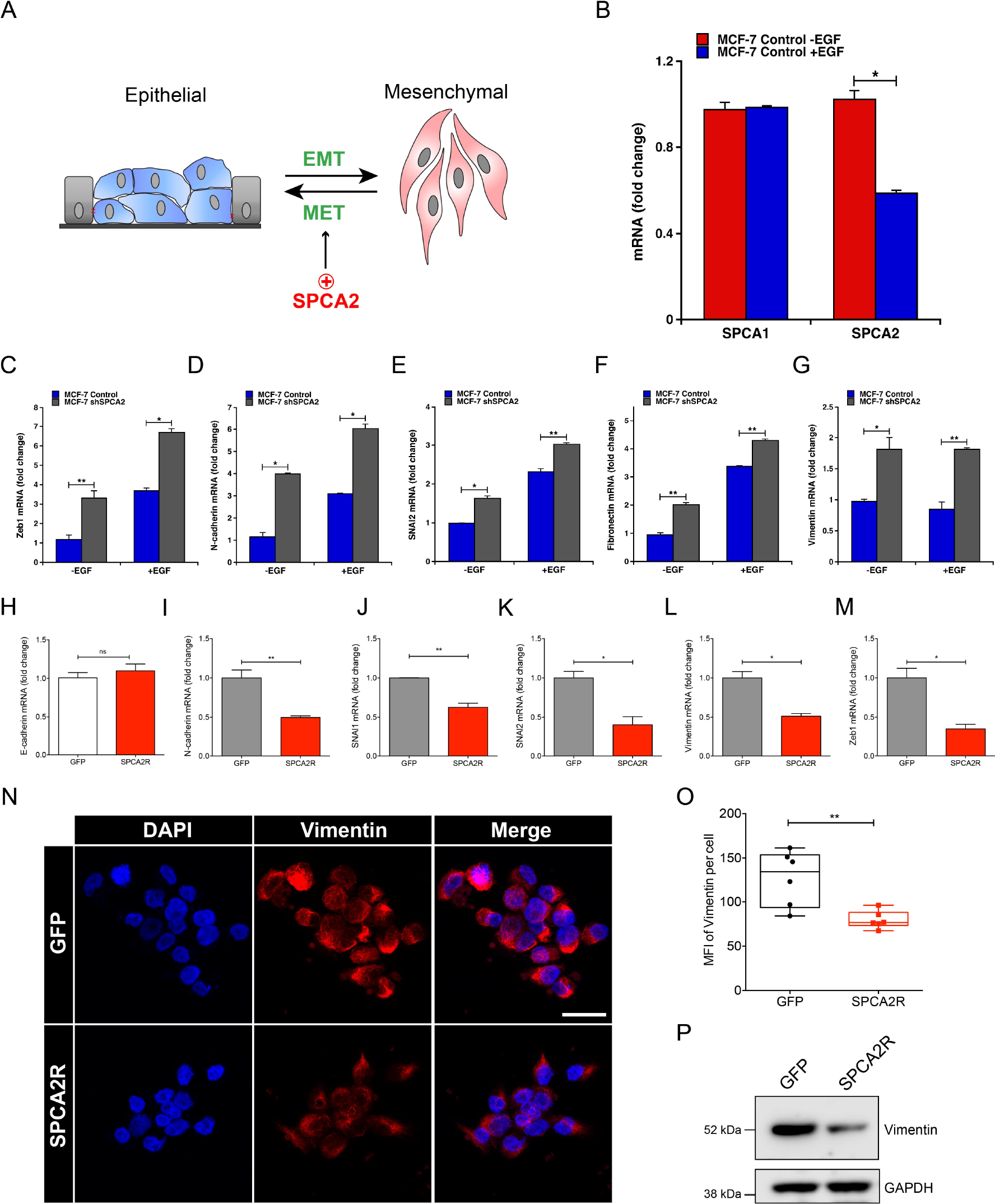
SPCA2 regulates gene expression in epithelial mesenchymal transition. (A) EMT-MET transitions may be regulated SPCA2 as shown (B) Effect of EGF (20 ng/ml) on mRNA levels of SPCA1 and SPCA2 in MCF7 cells determined by qPCR. *p<0.05, Student’s t-test. Gene expression changes accompanying EMT-MET transitions were evaluated following knockdown of SPCA2 in MCF7 (C-G) or upon expression of SPCA2R in MDA-MB-231 cells (H-M). MCF7 cells (scramble control or SPCA2 knockdown) were treated with 20 ng/ml EGF where indicated and transcript levels of the following mesenchymal markers were determined by qPCR: Zeb1 (C), N-cadherin (D), SNAI2 (E), fibronectin (F), and vimentin (G). *p<0.05, **p<0.01, Student’s t-test. Effect of SPCA2R expression in MDA-MB-231 cells, relative to vector control, on a panel of epithelial (H) and mesenchymal (I-M) gene markers was quantified by qPCR. n.s. not significant, *p<0.05, **p<0.01, Student’s t-test. (N) Representative confocal microscope images showing immunofluorescence staining of vimentin in MDA-MB-231 cells expressing SPCA2R, relative to vector control (40x magnification). (O) Vimentin mean fluorescence intensity (MFI) was quantified by ImageJ software from 6 images each condition. Each data point represents one image. **p<0.01 Student’s t-test, n=95 for GFP control and n=99 for SPCA2 overexpression. (P) Vimentin protein expression was detected using western blotting, GAPDH was used as a loading control. For each condition, 30 μg of protein lysates was loaded. See Fig. S5 and S6.

Next, we sought to determine if loss of SPCA2 is a driver of EMT, and conversely, if SPCA2 promotes MET (Fig. 6A). Strikingly, SPCA2 knockdown was sufficient to induce transcription of EMT-associated genes even in the absence of EGF, with further elevation of transcript in most genes upon EGF treatment (Fig. 6C-G; gray bars). In contrast, introduction of SPCA2R in MDA-MB-231 cells had the opposite effect of suppressing transcription of a range of mesenchymal markers, including N-cadherin, SNAI1, SNAI2/SLUG, Vimentin and ZEB1 (Fig. 6I-M), although not of E-cadherin (Fig. 6H) consistent with a post-translational role for SPCA2 in E-cadherin expression. Similar results were observed upon ectopic expression of SPCA2R in MCF10A and Hs578T (Fig. S6). We confirmed SPCA2R-mediated loss in expression of vimentin in MDA-MB-231 cells by confocal microscopy (Fig. 6N-O) and by Western blotting (Fig. 6P). These results demonstrate a cause and effect relationship between SPCA2 and EMT, pointing to a novel role for SPCA2 as driver rather than a passenger in the gene expression changes that accompany EMT-MET programs.

Finally, we investigated if gene expression changes elicited by SPCA2R in MDA-MB-231 cells were downstream of the Hippo signaling pathway. Addition of YAP inhibitor Verteporfin (40) to SPCA2R-transfected MDA-MB-231 cells rescued the expression of SNAI1, Vimentin, and ZEB1, key architects of EMT (Fig. S6 panels M-O). There was no significant rescue in the expression of CDH2 and SNAI2 (Fig. S6P-Q), indicating that additional or redundant pathways may be involved. These results show that the co-activator protein YAP plays a role in SPCA2-mediated EMT.

### SPCA2 suppresses cancer cell migration and tumor metastasis

Cell motility and migration underlie metastatic progression and are key features of EMT. We assessed the effects of SPCA2R expression on migration and invasion in the highly metastatic MDA-MB-231 cells, which exhibit very low endogenous levels of SPCA2 (Fig 3A). As shown in Figure 7A-B, wound-closure assays revealed that SPCA2R inhibited *in vitro* migration of MDA-MB-231 at all time points tested (p<0.05). Similarly, expression of SPCA2R inhibited cell migration in Boyden chambers by 40% (Fig S7A-C).

**Figure 7.**
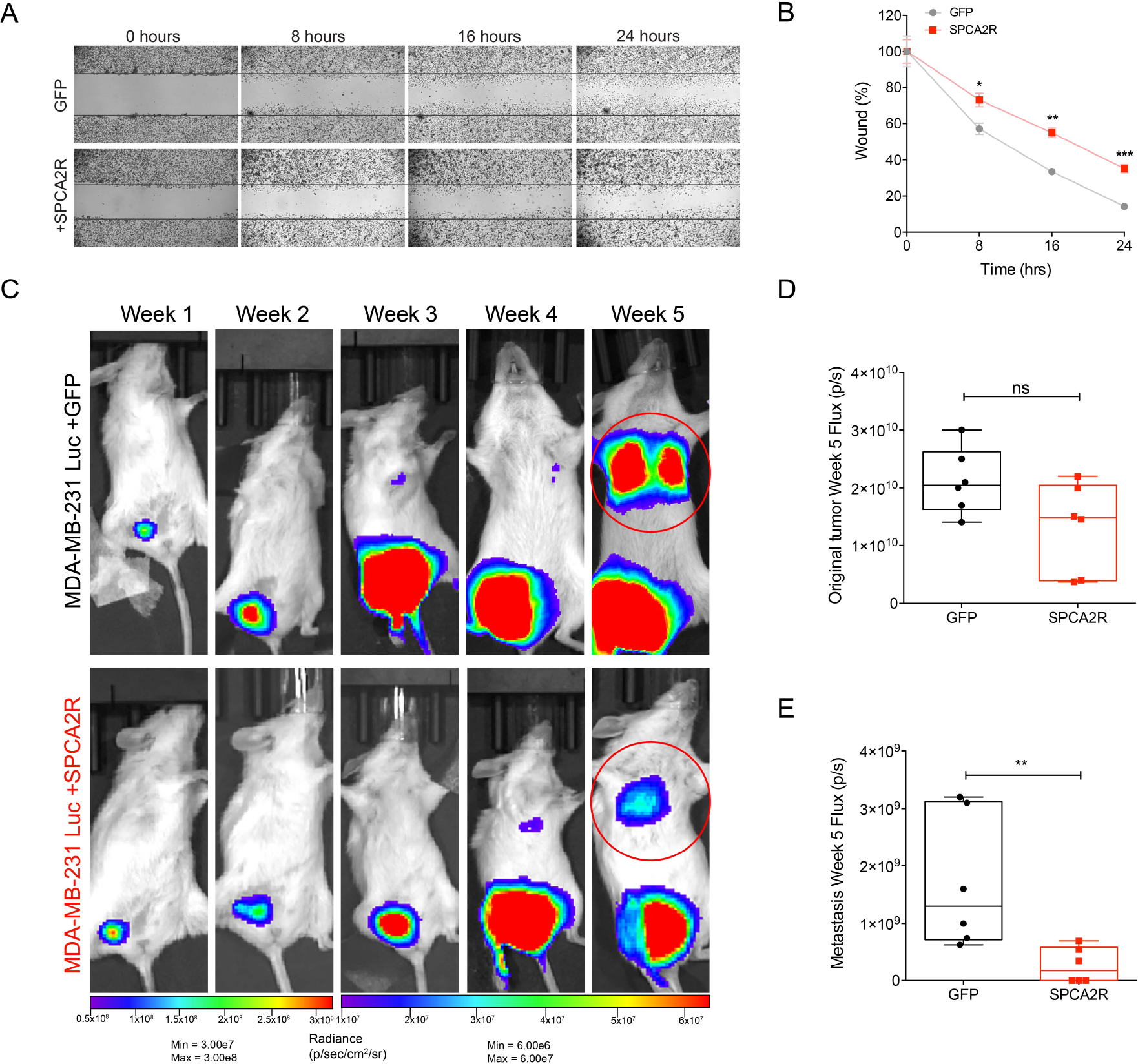
SPCA2 suppresses migration *in vitro* and metastasis *in vivo*. (A) Representative images of wound healing in MDA-MB-231 at the indicated times in control and SPCA2R transfected cells (2.5x magnification). (B) Quantification of cell migration expressed as percentage of control. SPCA2 significantly reduced migration at 8 h, 16 h, and 24 h. *p<0.05, **p<0.01, ***p<0.01. Student’s t-test, n=3 for each condition. (C) Bioluminescent images of NSG mice following engraftment with MDA-MB-231 Luc + GFP and MDA-MB-231 Luc + SPCA2R at indicated times and luminescence scales. (D-E) Quantification of bioluminescence in luciferase-labeled MDA-MB-231 Luc. Tumor growth and metastasis at Week 5 were represented as mean total flux (p/sec) α SEM. SPCA2 significantly reduced metastasis compared with control group. **p<0.01, Student’s t-test, n=6 for each condition. See Fig. S7.

Next, we investigated if SPCA2 has anti-metastatic effect *in vivo* using bioluminescent model MDA-MB-231 Luc. We injected MDA-MB-231 Luc + GFP and MDA-MB-231 Luc + SPCA2R cells in the mammary fat pad of NSG mice, and monitored metastasis using whole-body bioluminescence imaging at weekly intervals (Fig. 7C, Fig. S7D). Although there was no significant difference in the primary tumor size (Fig. 7D), tumors from cells transfected with SPCA2R showed significantly less metastasis compared to the control group at week 5 (Fig. 7E; p<0.01) with 3 of 6 mice showing a lack of lung metastasis. We suggest that the SPCA2-mediated suppression of mesenchymal genes demonstrated *in vitro*, (Fig 6, Fig. S6) confers inhibitory effects on tumor migration and metastasis.

## DISCUSSION

We show an isoform-specific post-translational role for SPCA2 in E-cadherin biogenesis that has functional consequences on cell adhesion and the epithelial-mesenchymal transition in the development of breast cancer metastasis. The specific molecular details of this mechanism will require further investigation and may involve a chaperone-like function of SPCA2 as previously described for Orai1 (20), isoform-specific differences in ion transport, Ca^2+^/Mn^2+^ selectivity or regulation that could alter the secretory pathway ionic milieu (41), or a new requirement for cytoplasmic Ca^2+^ signaling in E-cadherin trafficking or stability. It is unclear whether the secretory pathway plays a role in supplying Ca^2+^ to the extracellular/lumenal cadherin repeats known to mediate their structural rigidity and extended conformation (42). Interestingly, defective desmosomal adhesion is the underlying cause in Hailey-Hailey disease, an ulcerative skin disorder resulting from loss of one functional copy of the housekeeping isoform SPCA1 (43). Biogenesis of desmosomal components, including cadherins (desmoglein and desmocollin) and desmoplakins was delayed upon knockdown of SPCA1 in keratinocytes (30). In light of our new data showing that loss of SPCA2 leads to defective biogenesis of E-cadherin, a component of the adherens junction, we propose that both secretory pathway Ca^2+^-ATPase isoforms play similar but non-redundant roles in the establishment of cell adhesion.

Given the developmental and pathological implications of the EMT program, intensive efforts have been directed toward understanding the underlying molecular mechanisms and regulatory networks. Down-regulation of E-cadherin is a defining mechanism in EMT. Most studies in the literature have focused primarily on silencing mutations in E-cadherin and transcriptional repression (3). However, a significant percentage of invasive tumors have normal E-cadherin gene and transcription, which may point to post-translational down-regulation of E-cadherin, although much less is known about these mechanisms (44). Here, we show that loss of SPCA2 essentially phenocopies loss of E-cadherin in tumorsphere formation, Hippo pathway signaling, and in gene regulation and tumor phenotypes associated with epithelial mesenchymal transition. Previously, dysregulation of membrane trafficking of E-cadherin has been implicated in EMT associated with viral oncogenesis, such as Rous sarcoma and Kaposi’s sarcoma (45, 46). This study showing SPCA2-mediated regulation of E-cadherin expression offers a new mechanism to regulate adherens junction stability in non-viral oncogenesis.

YAP/TAZ are essential for initiation and growth of most solid tumors and their activation is critical for cancer stem cell attributes and metastasis (47). Importantly, YAP is a sensor of the structural and mechanical cues affecting tumor microenvironment and mediates contact inhibition downstream from E-cadherin (4, 8). The Hippo/YAP pathway presents a therapeutic vulnerability of cancer cells that can be exploited. Thus, our finding that SPCA2 is an upstream regulator of this pathway is an important advance that provides a mechanistic explanation for the role of SPCA2 in EMT-MET transitions via down regulation of E-cadherin. Furthermore, store-independent Ca^2+^ entry (SICE) mediated by SPCA2 demonstrates, for the first time, a role for Ca^2+^ in YAP phosphorylation in breast cancer cells that complements recent observations on YAP phosphorylation by store-operated Ca^2+^ entry (SOCE) in glioblastoma cells (35) and provides additional avenues for therapeutic intervention.

Down-regulation of SPCA2 is associated with basal and claudin-low breast cancer subtypes (19) that are characterized by hormone receptor negativity, and a high degree of invasion and metastasis. Therefore, low SPCA2 may be a useful prognostic marker of the malignant invasive phenotype. In contrast, high SPCA2 is characteristic of luminal A/B and HER2+ subtypes where it contributes to the formation of breast microcalcifications associated with tumorigenesis (19). We propose a model to summarize the role of SPCA2 in breast cancer. During the early stages of breast oncogenesis, proliferation and dysplasia of epithelial cells results in carcinoma *in situ*. Tumor cells in this stage express high levels of SPCA2, elevated cytoplasmic Ca^2+^ and rapid cell proliferation (16). Biogenesis and trafficking of E-cadherin is normal, cell-cell contacts are maintained and EMT genes are repressed. Induction of EMT in a subset of breast cancer cells is associated with down-regulation of SPCA2 and other ESG, resulting in disrupted E-cadherin trafficking and enhanced mesenchymal gene expression that facilitates intravasation of tumor cells and escape into the blood stream. Subsequently, extravasation of a subset of these breast cancer cells from the blood stream into tissues could be followed by mesenchymal to epithelial transition (MET), which reinstates expression of SPCA2 and other epithelial genes such as E-cadherin (48) leading to secondary tumors at metastatic sites.

## Author Contributions

**Conception and Design**: D.K. Dang, M. R. Makena, J.P. Llongueras, H. Prasad, R. Rao

**Development of Methodology**: D.K. Dang, M.R. Makena, J.P. Llongueras, H. Prasad

**Acquisition of data**: D.K. Dang, M. R. Makena, J.P. Llongueras, M. Bandral

**Analysis and interpretation of data**: D.K. Dang, M.R. Makena, J.P. Llongueras, H. Prasad, M.J. Ko, R. Rao

**Writing, review and revision of manuscript**: D.K. Dang, M.R. Makena, J.P. Llongueras, H. Prasad, R. Rao

**Administrative, technical or material support**: M. Bandral, M.J. Ko

**Study supervision**: R. Rao

## Acknowledgements

This work was supported by a grant from the National Institutes of Health (DK103078) to R.R. H.P. was a Fulbright Fellow supported by the International Fulbright Science and Technology Award, J.P. was supported in part by the NIH PREP scholar program at Johns Hopkins and M.B. was supported by the Khorana Program for Scholars.

