## Supplemental Figures S1-S7 for "A Ca^2+^-ATPase Regulates E-cadherin Biogenesis and Epithelial-Mesenchymal Transition in Breast Cancer Cells"

Figure S1

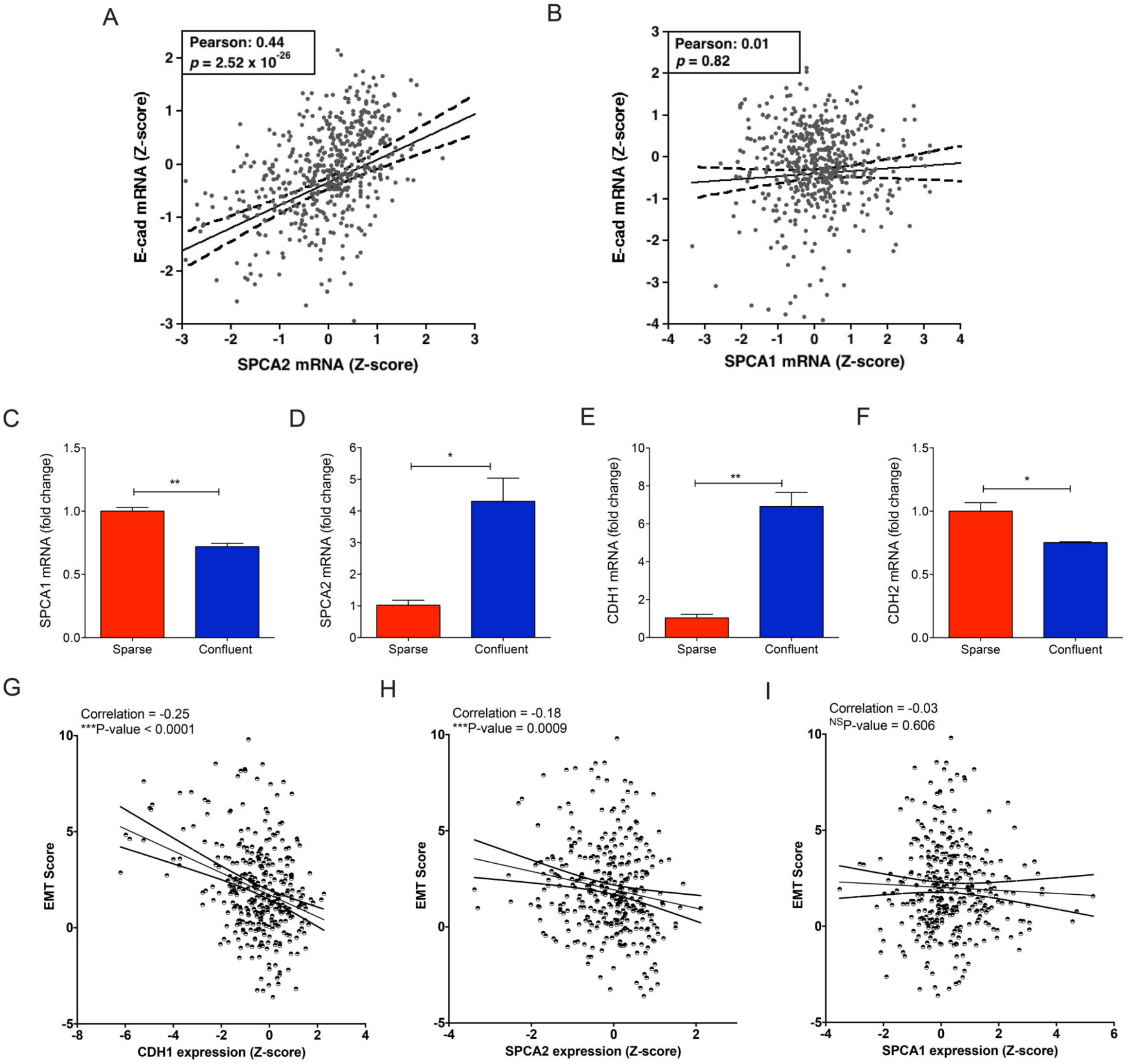


**Figure S1. SPCA2 is transcriptionally linked with E-cadherin in breast cancer**

(A) Correlation of E-cadherin and SPCA2 mRNA expression in cancer cell lines. Pearson coefficient = 0.44, $p=2.52\times10^{-26}$. (B) mRNA levels of SPCA1 do not correlate with E-cadherin mRNA. p=0.82; Pearson coefficient = 0.01. (C-F) qPCR analysis of SPCA1, SPCA2, E-cadherin (*CDH1*) and N-cadherin (*CDH2*) gene expression in cDNA extracted from MDA-MB-231 cells cultured under sparse and confluent conditions. **p<0.01, *p<0.05, t-test, n=3. (G-I) Analysis of 633 breast tumors from TCGA shows EMT score is negatively correlated with expression of E-cadherin (*CDH1*, corr. -0.25, p<0.0001) and SPCA2 (corr. -0.18, p=0.0009), but not with SPCA1 (corr. -0.03, p=0.606).

#### Figure S2. Selective effects of SPCA gene knockdown on protein expression

(A) Knockdown of SPCA2 in MCF-10A cells was confirmed by qPCR. **$p<0.01$, n=3, Student's t-test. (B) qPCR analysis showed no change in E-cadherin (*CDH1*) transcript. n.s. not significant, n=3, Student's t-test. (C) Western blot of cell lysates probed with antibody against E-cadherin and GAPDH. (D) E-cadherin protein expression from densitometric analysis of Western blots, normalized against GAPDH. *$p<0.05$, Student's t-test, n=3. (E-F) Western blotting of E-cadherin in lysates from MCF7 cells treated with control or shSPCA1; protein levels were normalized to β-Actin and quantified from three independent transfections. (G-H) Western blotting of EGF receptor in MCF7 cells treated with control or shSPCA2; protein levels were normalized to GAPDH and quantified from three independent transfections. (I-J) Western blotting of ZO-1 in MCF7 cells treated with control or shSPCA2; protein levels were normalized to GAPDH and quantified from three independent transfections.

Figure S2

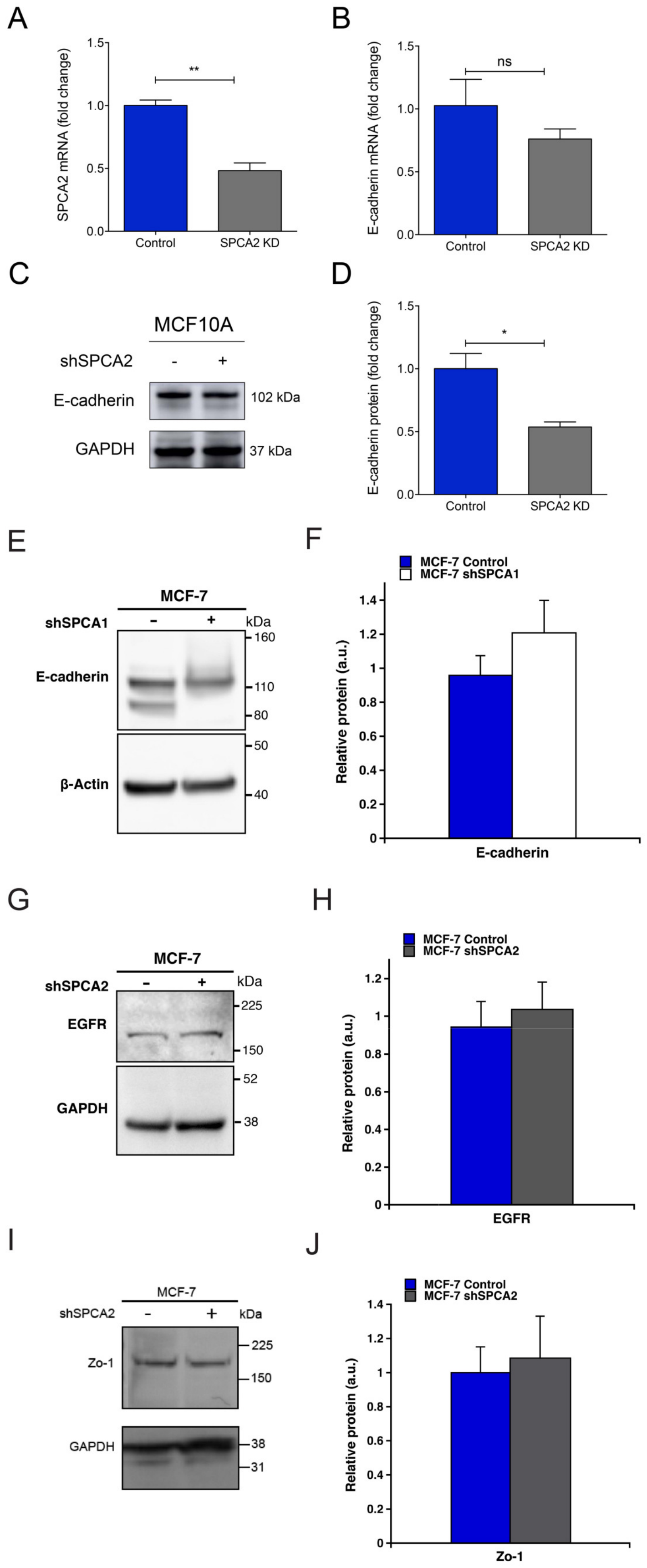
A
SPCA2 mRNA (fold change)
1.5
1.0
0.5
0.0
**
Control
SPCA2 KD
B
E-cadherin mRNA (fold change)
ns
Control
SPCA2 KD
C
MCF10A
shSPCA2
-
+
E-cadherin
102 kDa
GAPDH
37 kDa
D
E-cadherin protein (fold change)
*
Control
SPCA2 KD
E
MCF-7
shSPCA1
kDa
160
110
80
50
40
E-cadherin
β-Actin
F
MCF-7 Control
MCF-7 shSPCA1
Relative protein (a.u.)
E-cadherin
G
MCF-7
shSPCA2
EGFR
GAPDH
225
150
52
38
H
MCF-7 Control
MCF-7 shSPCA2
EGFR
I
MCF-7
Zo-1
GAPDH
31
J
Zo-1

### Figure S3

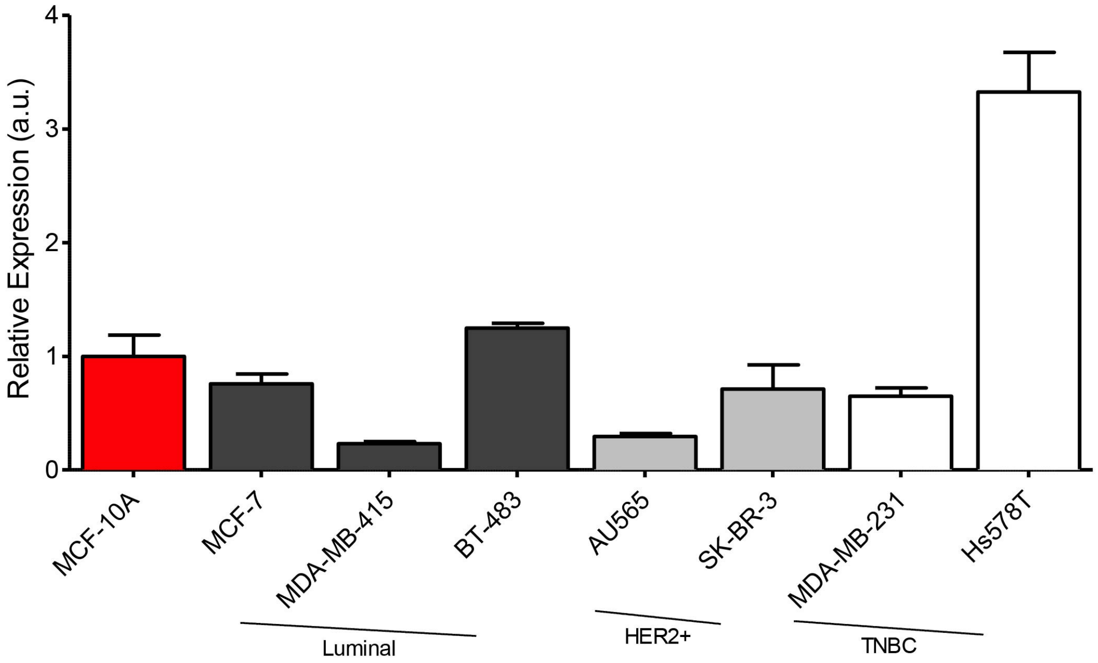


**Figure S3. SPCA1 expression in breast cancer cell lines**

A panel of breast cancer cell lines classified as luminal, HER2+ or TNBC subtypes as indicated, was evaluated for SPCA1 expression by qPCR. Results are normalized to MCF-10A.

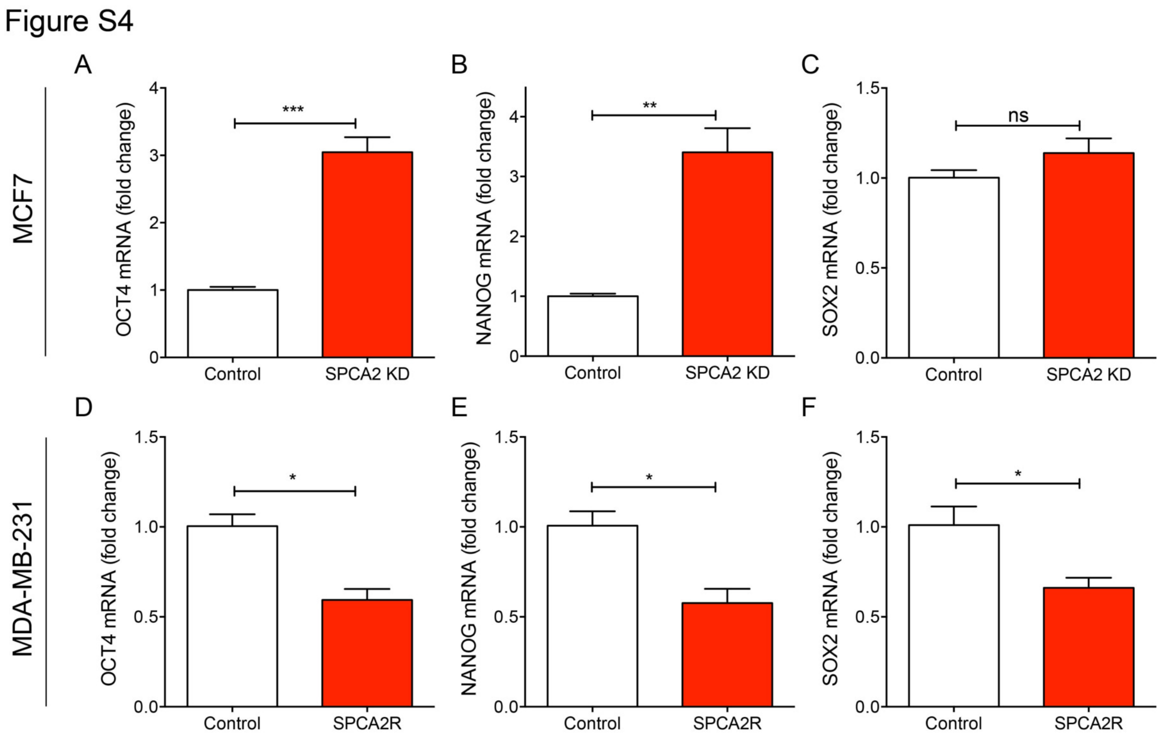


**Figure S4. SPCA2 alters expression of cancer stem cell gene markers**

(A-C) Effect of SPCA2 knockdown in MCF7 cells, and (D-F) SPCA2R expression in MDA-MB-231 cells, relative to control, on stem cell gene markers, quantified by qPCR. n.s. not significant, *p<0.05, **p<0.01, ***p<0.001Student's t-test.

Figure S5

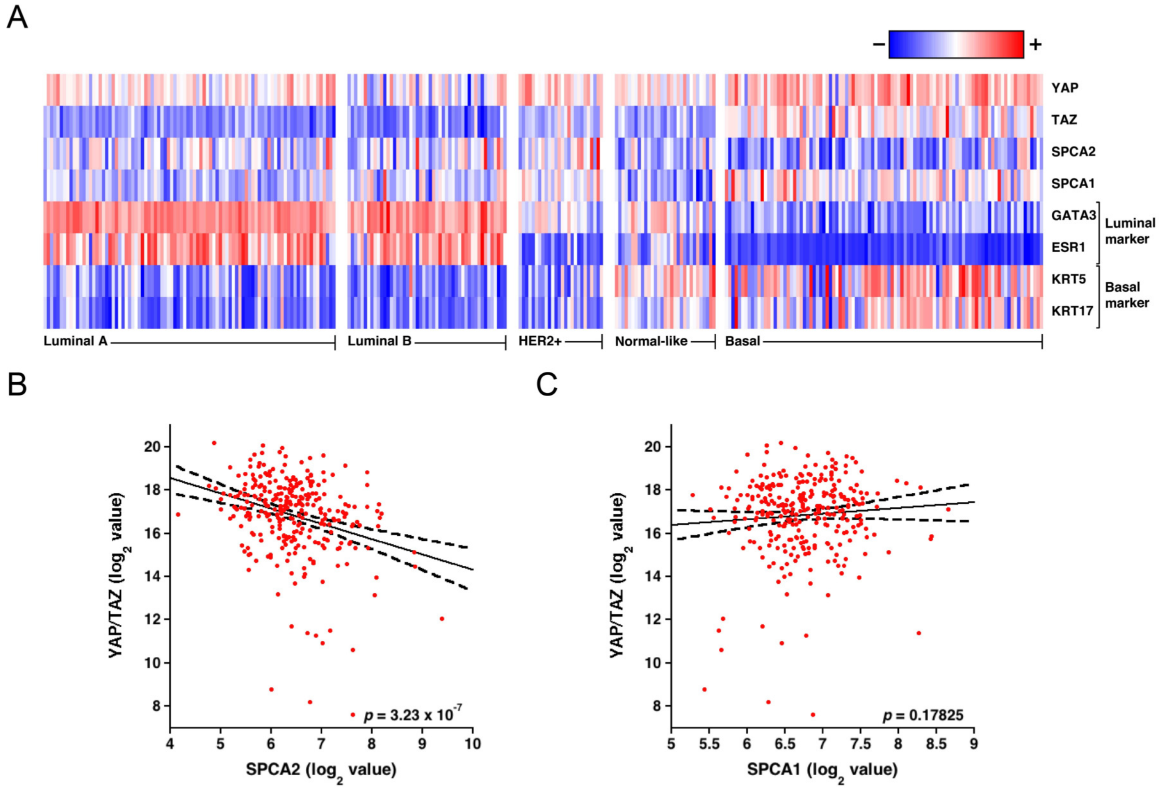


**Fig. S5. Transcriptional links between EMT-associated genes and SPCA**

(A) Expression heatmap of YAP, the related co-activator TAZ, SPCA1 and SPCA2 in patient breast cancer tissue obtained from the GSE31448 microarray dataset. Note the reciprocal expression of SPCA2 with YAP and TAZ, particularly in basal-like breast cancer. (B) Scatter plot of SPCA2, and (C) SPCA1 expression against YAP/TAZ expression levels from all breast cancer subtypes (n=295). SPCA2, but not SPCA1, is negatively correlated with YAP/TAZ expression ($p=3.23X10^{-7}$).

Figure S6

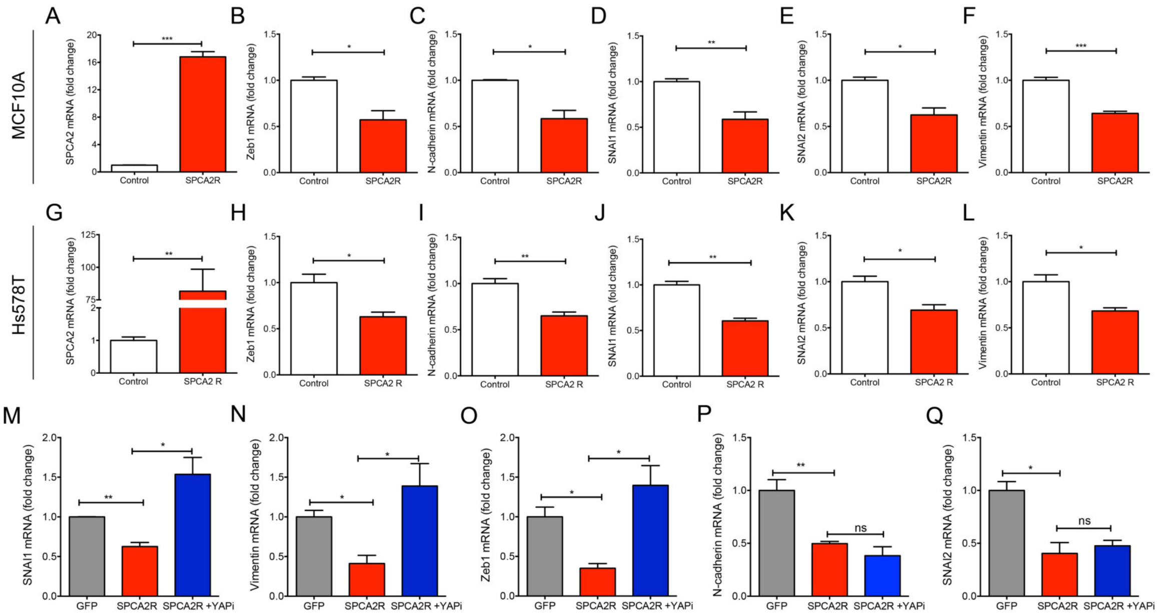


**Figure S6. SPCA2 suppresses mesenchymal markers in multiple breast cancer cell lines**

(A-F) Ectopic expression of SPCA2 in the non-tumorigenic mammary epithelial cell line MCF10A significantly suppressed mesenchymal gene markers *CDH2*, *SNAI1*, *SNAI2*, Vimentin and Zeb1. *p<0.05, **p<0.01, Student's t-test, n=3 for each condition. (G-K) Ectopic expression of SPCA2 in TNBC cell line HS578T significantly suppressed mesenchymal gene markers *CDH2*, *SNAI1*, *SNAI2*, Vimentin and Zeb1. *p<0.05, **p<0.01, Student's t-test, n=3 for each condition. (M-Q) Effect of YAP inhibitor Verteporfin (10 μM for 24 hours) on SPCA2R induced EMT gene markers in MDA-MB-231 cells, relative to vector control, quantified by qPCR. n.s. not significant, *p<0.05, **p<0.01, Student's t-test, n=3.

### Figure S7

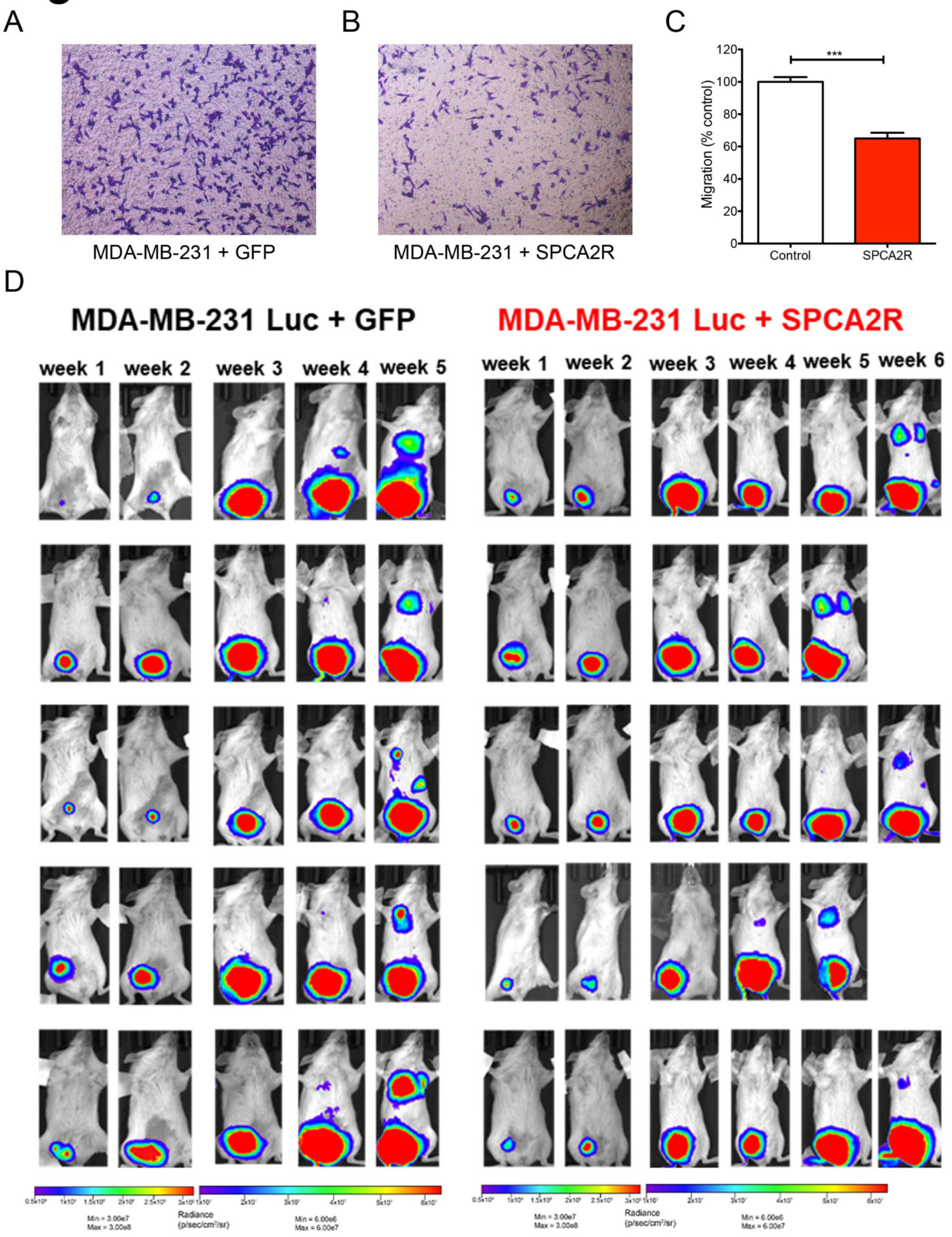


**Figure S7. SPCA2 suppresses metastasis of MDA-MB-231 cells *in vivo***

(A-B) Representative microscope images of MDA-MB-231 cells transfected with vector control or SPCA2R, in Boyden chamber (10x magnification). (C) Migration was quantified by ImageJ software from 5 images each condition. ***p<0.001 Student's t-testBioluminescent images of NSG mice engrafted with MDA-MB-231 Luc + GFP and MDA-MB-231 Luc + SPCA2R tracking luciferase activity at different time points. Week 1 and 2 images were represented using bioluminescent intensity; maximum (max) = $3.0x10^8$; minimum (min) = $3.0x10^7$. To illustrate metastasis, week 3, 4 and 5 images were represented using bioluminescent intensity; min = $6.0x10^6$, max= $6.0x10^7$.
